# Mapping a viral phylogeny onto outbreak trees to improve host transmission inference

**DOI:** 10.1101/010389

**Authors:** Jonathan E Allen, Stephan P Velsko

**Affiliations:** Lawrence Livermore National Laboratory, Livermore, CA, 94551, USA

## Abstract

**Background:** Developing methods to reconstruct transmission histories for viral outbreaks could provide critical information to support locating sources of disease transmission. Phylogenetic methods used to measure the degree of relatedness among sequenced viral samples have proven useful in identifying potential outbreak sources. The complex nature of infectious disease, however, makes it difficult to assign a rigorously defined quantitative confidence value assessing the likelihood of a true direct transmission event using genetic data alone.

**Results:** A new method is presented to calculate a confidence value assessing the likelihood of a transmission event using both phylogenetic inference and limited knowledge of incubation and infectious duration times. The method is applied to simulations of a foot and mouth disease (FMD) outbreak to demonstrate how the combination of both phylogenetic and epidemiology data can be used to strengthen the assessment of the likelihood of direct transmission over methods using just phylogenetic data or infection timing data alone. The method is applied to a previous FMD outbreak to identify areas where over confidence in previously inferred direct transmission may exist.

**Conclusion:** Combining knowledge from viral evolution and epidemiology within a single integrated transmission inference framework is an important approach to assess the potential likelihood of transmission events and makes clear how specific features of a virus’ spread through the course of an outbreak will directly determine the potential for confidence in inferred host transmission links.

## Background

Developing methods to protect against RNA virus outbreaks in the face of constantly changing disease dynamics remains one of the great public health challenges of our time. One small piece of this problem involves determining the host-pathogen transmission history to identify an outbreak’s origin. In recent cases such as the 2003 SARS outbreak [1, 2], H5N1 influenza [3] infection in humans and Marburgvirus [4], phylogenetic inference has been used to help identify potential outbreak sources. Typically, a collection of sequences from candidate related sources are compared to infer transmission links between the most closely related sequences. These methods have been used beyond the public health domain to provide evidence in HIV [5, 6] and other forensic investigations [7] to help infer when a suspect is involved in infecting a victim.

**Figure 1:**
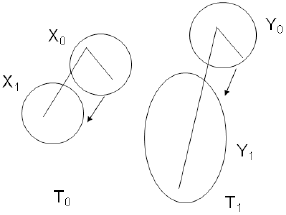
Schematic of transmission links overlaid on phylogenetic trees *T*_0_ (left tree) and *T*_1_ (right tree). Transmission links are shown as directed edges connecting two infected hosts *X*_0_ to *X*_1_ and *Y*_0_ to *Y*_1_ respectively.

While phylogenetic data helps establish the evolutionary relationships among the sequenced viral samples, there are no well established tools to translate these results to explicitly infer host to host transmission links with confidence values that accurately capture the uncertainty of host transmission inference. The result is a sampling and transmission inference process that could lead to false confidence when using genetic data to establish host to host transmission [8, 9].

Complementary to the viral genome sequencing advances is contact tracing work to find host to host transmission patterns independent of molecular genetic sequence data [10–13]. Previously a transmission inference framework was described that estimates the probability of viral transmission between two hosts using both viral sequence samples and contact tracing data [14, 15]. The method uses labeled training sets from past outbreaks to correlate models of viral evolution associated with direct transmission and indirect transmission to assign a posterior probability that gives a confidence value for inferring a transmission relationship between two infected hosts. One of the limitations of the approach is the need for a robust training set, which can limit application to novel outbreaks. Even when training data is available, a training set over-represented with local outbreak features could skew confidence values. Thus, there is a need to develop additional tools to mitigate limits introduced from training. One feature that can be difficult to accurately account for with limited exemplars is time series data. The objective of this article is to present an approach to incorporate time series data with molecular genetic sequence data to measure the uncertainty of host to host transmission events in the face of limited training data and variability in host infection times.

Figure 1 shows a simple schematic to illustrate the variable host infection time transmission inference problem. The figure shows two simple trees, *T*_0_ and *T*_1_ (left and right tree respectively) where the infected hosts are overlaid as circles on the trees to show the transmission event from host *X*_0_ to *X*_1_ in *T*_0_ and *Y*_0_ to *Y*_1_ in *T*_1_. The larger circle for *Y*_1_ represents a longer infection time and the two trees represent the evolutionary relationship between the sequences. Assuming comparable homogeneous nucleotide substitution rates, the longer branch length in *T*_1_ could be explained by the longer infection time in host *Y*_1_ prior to sampling. Including the direct transmission branch lengths from *T*_0_ and *T*_1_ in a single genetic based transmission inference model leads to a distribution of genetic distances associated with direct transmission for a wider range of values. Thus, estimating accurate confidence values means ensuring that a representative sampling from the true distribution of infection times can be obtained. To measure the impact of this problem, a new method is introduced to incorporate variation in infection times when assigning a final posterior probability confidence value. The proposed method forgoes the use of training data by relying on prior knowledge of viral infection times to act as a constraint in conjunction with phylogenetic data. The results show that with knowledge of infection times confidence in transmission inference can be high and even with limited knowledge of timing patterns, high confidence values can be obtained that exceed the training based methods. Thus, this new transmission inference approach allows for the use of clinical observations of infection to be combined with inferred viral evolution when training data is not available to measure uncertainty associated with specific transmission hypotheses.

Efforts to integrate models of epidemiology and genetic evolution have been ongoing for some time [16–19], yet less work has focused on the specific problem of measuring the certainty of inferring individual host transmission events. One related effort is a method called SeqTrack, which was used to examine the origins of the 2009 H1N1 outbreak [20]. This work is similar to our previous transmission inference [14] approach with a few key differences. Rather than inferring a host transmission graph, a genealogy tree is inferred where nodes represent sequence samples and edges between nodes link a sequence to putative descendants. Sample collection times exclude some candidate edges and edge weights represent genetic distances between two sequences. The genealogy tree is inferred by finding the directed graph analog of the minimum spanning tree. Given the elapsed time between two sequences and a per day mutation rate, a confidence value is assigned for direct genealogy linkages. The framework presented here is a more general framework where variable within host evolution times are assumed to play a role. A second closely related effort is the work to recover transmission links between farms applied to the 2001 FMD outbreak [21].

FMD samples were collected and sequenced and an inferred phylogenetic tree was used in conjunction with farm infection times to evaluate the most likely transmission tree. Data from that study is used here and a comparison with this work will be covered in detail later in the paper. While the Cottam et al. approach evaluates a putative outbreak tree overlaid onto a single phylogenetic tree, our main contribution is to provide a framework that models features of transmission inference not previously considered. In particular, uncertainty in transmission inference is newly captured by explicitly considering unobserved nodes in the outbreak tree and uncertainty of phylogeny through the use of multiple potentially competing phylogenetic inferences. The result is a more realistic measure of uncertainty when assigning a transmission link. The framework’s added use of competing transmission hypotheses does, however, come with an increased computational cost. The remainder of the paper details the modifications to our original framework, shows how new information is used in simulations and real data, details important differences with the Cottam et al. approach and introduces a way for dealing with limitations introduced by the framework’s increased computational cost.

## Results

### Algorithm

The previous framework is briefly reviewed to motivate the notation used to incorporate outbreak timing data. For any two infected hosts A and B, distance *δ* = *δ*(*A, B*) defines a genetic distance between samples taken from the two hosts. In the simplest form one sequenced sample is available for each host and the genetic relationship is described by a Jukes-Cantor like distance measure (*δ* can take on more complex forms as will be described later). The variable M represents the number of edges (ignoring direction) linking two nodes in an outbreak tree where nodes represent infected entities and edges represent directed transmission events from a source to a recipient. For the direct transmission hypothesis M=1 the posterior probability is defined to be:

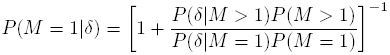

where 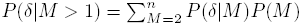 for some maximum transmission chain length *n*. *P* (*δ|M)* is estimated from labeled training data using density estimators or fitting the observed data to a distribution such as Poisson, Negative Binomial or Gamma.

The modified framework explicitly combines knowledge of viral infection times within the genetic based transmission inference framework. This is accomplished by adding a parameter *σ* and computing the value *P* (*δ, σ|M)* rather than *P* (*δ|M)*. This reflects the desire to estimate the probability of observing the genetic relation *δ* in conjunction with infection related timing values for the transmission hypothesis *M*. The *σ* parameter values are motivated by a SEIR type infection concept [22], which describes a host in one of four states 1) Susceptible (S) to infection 2) Exposed (E) to an infection (but not infectious) 3) Infectious (I) - able to transmit virus to other hosts and 4) Recovered state (R) - where the host is no longer able to transmit the virus. The SEIR model describes outbreak dynamics in a homogeneous mixing population and since the focus of the framework is to predict individual host to host transmission relationships a pre-determined outbreak tree is used, which excludes hosts in the susceptible state. Figure 2 shows the graphical representation of two infected hosts A and B linked by a direct transmission event. Each circle represents a discrete time point marking the transition between states or time intervals. Transmission time points marked red in Figure 2 denote the time of transmission from Host A to Host B. Sample time points marked in purple represent times when viral samples are collected for sequencing. The representation is analogous to a phylogenetic tree where branches (or tree edges) represent time intervals and nodes represent fixed time points. Here edges or transmission events between infected nodes represent instantaneous time events and the host nodes represent time intervals defined by a linear chain of discrete events. The motivation for this representation is to employ a sufficiently realistic epidemiology model to describe the transmission relationships of interest and use existing methods for modeling viral evolution by mapping them to the epidemiology model.

**Figure 2:**
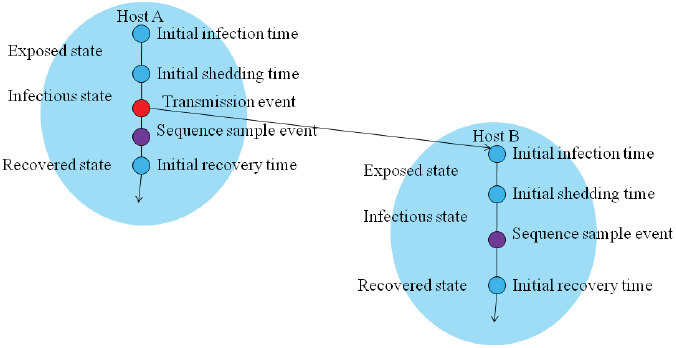
Host infection time model. The small circle nodes denote discrete time events and edges denote time intervals. The large blue circles denote a single infected host.

The modified framework reports the probability of jointly observing both the inferred genetic relationship between sequences from host A and B and the timing events associated with the viral infections of the two hosts. A key phylogenetic inferred parameter becomes the estimated “wall clock” time of the inferred most recent common ancestor between the time stamped sequence samples taken from the two hosts and potentially using samples from the larger outbreak. Here the Bayesian estimation software BEAST [23] is used to estimate a series of candidate phylogenetic relationships and inferred times, weighted by a likelihood value for each estimate. Transmission hypothesis testing determines whether the constraints implied by the hypothesis are consistent with the inferred most recent common ancestor times. Next, a description for estimating the probability of direct transmission is given, which is then extended to evaluate any transmission relationship between two nodes in an outbreak tree.

Probability distributions describe the expected exposure time and the expected infectious time in the host. An additional key requirement is the assignment of a fixed time point to each observed host that is relative to some global time clock. The point in time context can always be satisfied by assuming that the time is noted when a sample is collected from each infected individual. An alternative is to assume a fixed recovery time. For notational simplicity, a fixed recovery time is used although a comparable procedure is available for working from sample times as the reference. With host X recovery time *R*_*X*_, infectious time duration *inf*_*X*_ occurring with probability *P* (*inf*_*X*_) and exposure time *exp*_*X*_ occurring with probability *P* (*exp*_*X*_), the direct transmission hypothesis that host *A* transmitted the virus to host *B* is calculated by evaluating all pairwise combinations of *exp*_*x*_ and *inf*_*x*_ for each host. Candidate transmission events are marked by cases where the infectious time interval for host A overlaps with a potential initial infection time for host B and the inferred most recent common ancestor time for the sequence samples occurs after the initial infection time of A and prior to the initial infection time of B.

Figure 3 shows the notation and the constraints for the most recent common ancestor time (MRCAT). The initial infection time *E*_*B*_ of host B is *E*_*B*_ = *R*_*B*_ − *inf*_*B*_ − *exp*_*B*_ and similarly the initial infection time *E*_*A*_ for host A is *E*_*A*_ = *R*_*A*_ − *inf*_*A*_ − *exp*_*A*_, which leads to the constraints defined by the specific transmission hypothesis. The transmission constraint *C*_1_(*E*_*B*_, *R*_*A*_, *I*_*A*_) returns 1 if *I*_*A*_ ≤ *E*_*B*_ < *R*_*A*_ and 0 otherwise. The most recent common ancestor constraint *C*_2_(*M RCAT, E_B_, E_A_*) returns 1 if *E*_*A*_ ≤ *M RCAT* < *E*_*B*_ and 0 otherwise. The probability of *E*_*B*_ and *E*_*A*_ is defined to be *P* (*E*_*B*_|*exp*_*B*_, *inf*_*B*_, *R*_*B*_) = *P* (*exp*_*B*_) *× P* (*inf*_*B*_) and *P* (*E*_*A*_|*exp*_*A*_, *inf*_*A*_, *R*_*A*_) = *P* (*exp*_*A*_) *× P* (*inf*_*A*_) respectively. Now the genetic relation parameter *δ* is a set of estimates *M RCAT_i_* for 1 *≤ i ≤ q* where *q* is a finite integer determined during an independent phylogenetic parameter estimation process. (For example, using Bayesian genetic inference software BEAST, a default value could be *q* = 10, 000 estimates.) The direct transmission hypothesis is tested by finding the infection time intervals that are consistent with the inferred most recent common ancestor times and is described as follows:

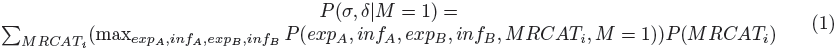

and

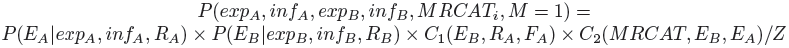

where Z is a normalizing constant that reflects the maximum probability value of the individual timing interval events independent of constraints.

**Figure 3:**
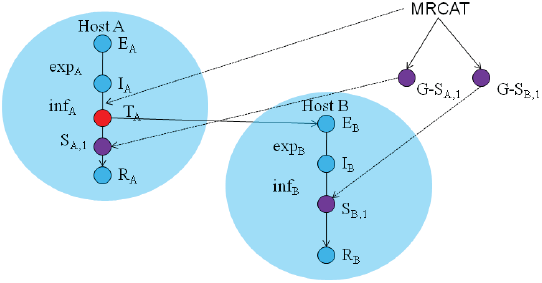
Example mapping of a two leaf phylogenetic tree onto the host node n (A or B). *E*_*n*_ is the initial infection time, *I*_*n*_ is initial infectious time, *T*_*A*_ is transmission time from host A to host B. *S*_*n*,1_ is the sample collection time (for the first of potentially multiple samples) and *R*_*n*_ is the initial recovery time. *G* – *S*_*A*, 1_ and *G − S*_*B*,1_ are genetic sequences for the sampled isolates. MRCAT is an inferred time that occurs within the time interval: *E*_*A*_ ≤ *M RCAT* < *T*_*A*_.

To maintain consistency with previous descriptions of the original framework [14], the transmission hypothesis *M* up to now was described by an integer value, which reports the number of edges in a tree separating the two nodes in question. This is sufficient for genetic only transmission inference when genetic distance is computed without regard to transmission direction. Thus, different sub tree configurations containing the same number of edges are treated identically without regard to transmission direction.

Figure 4 shows how the time series process of outbreak events distinguishes transmission direction. For a sub tree connecting two nodes with k edges, there are k distinct sub trees with each sub tree representing a distinct potential source node for the infection. For example, the transmission sub tree with two edges separating host A and B has two possible sub trees to describe the transmission relationship between the two hosts, one where the source node is A and one sub tree where A and B receive the infection through direct transmission from a common source. In general for a transmission hypothesis linking two nodes by k edges, there are k distinct sub trees that must be evaluated.

**Figure 4:**
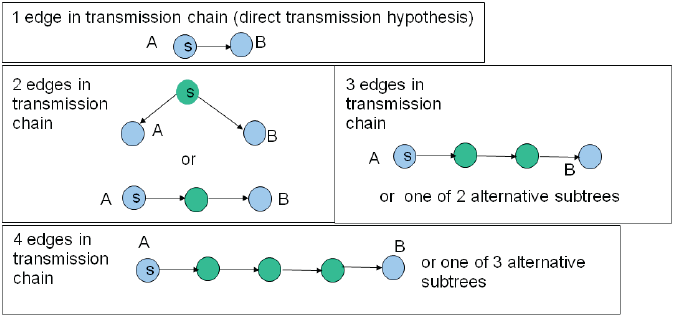
Transmission chains. The nodes labeled “s” represent the putative infection source. Blue nodes represent the two hosts (A and B) with sequenced samples. Green nodes reflect putative infected (hidden) intermediate hosts. Examples of transmission chains from length 1 (the direct transmission case) to 4 are shown.

The color coding of nodes in Figure 4 highlights the two distinct node types in the hypothesis testing framework. The hypothesis test considers an outbreak tree with “observed” nodes and “hidden” nodes. The hidden nodes are marked green in the figure and represent infected individuals where no direct observable data is available. The hidden nodes explicitly model the probability of the alternative transmission hypothesis linking two observed nodes A and B. To evaluate the *M >* 1 case an evaluation procedure for each sub tree *t*_*M*_ ∊ *T*_*M*_ (*A, B*) is used. Thus, *P* (*σ*, *δ|M* > 1) becomes the value of the most probable alternative hypothesis:

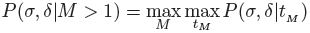

and the calculation of *P* (*σ, δ|tM)* invokes a recursive tree traversal procedure to check the possible transmission timing values that are consistent with the transmission hypothesis *t*_*M*_ . Hidden nodes cannot be tied to a global reference time but rather occur within a time range that is relative to the reference time points specified by the observed nodes and satisfy the transmission and phylogenetic constraints. A tree traversal process sets the range of possible infection times using the reference time points for the observed nodes. One of two traversal procedures is invoked depending on the sub tree structure: directed linear chain, where an observed node A is the source and the last node in the transmission chain is the other observed node B (see for example the bottom panel in Figure 4). The alternative traversal process is invoked when the two observed nodes are linked by transmission from a common source as shown in Figure 5. Figure 5 shows an example of the tree traversal process for one specific instance of *t*_*M*_ where *M* = 3. The common source sub tree traversal algorithm starts with each observed node (leaf) independently and moves up the tree to the root. Each node *n* is assigned an earliest infection time *E*_*min,n*_ and a latest infection time *E*_*max,n*_. The parent of node n is assigned an earliest and latest infection time for each minimum and maximum future initial infection time at node n. The total number of possible initial infection time ranges for the source node is 2*^d^/*2 where *d* is the length of the chain from the source to the leaf. The set of initial timing event ranges derived from one observable node are compared with the set of infection time ranges derived from the other observable node to determine whether there is a candidate infection time interval that supports the tree’s transmission hypothesis linking both observed nodes.

**Figure 5:**
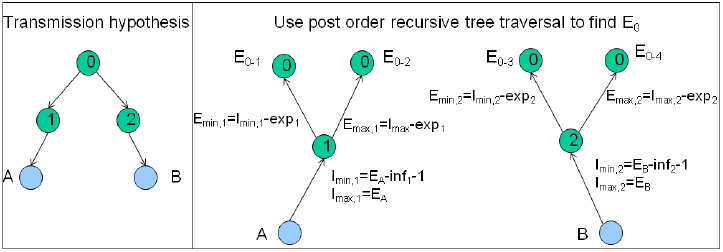
Tree traversal procedure (right panel) for the transmission hypothesis shown in the left panel. Left panel shows a transmission hypothesis linking two observed infected nodes A and B (marked in blue), connected by a common unobserved labeled source node marked 0 and two intermediate source nodes marked 1 and 2. The right panel shows the post order tree traversal, where the observed time stamp for each observed node serves as the starting point and the possible infection times for each source node are computed by moving up the tree.

The consistency function *g*(*t*_*M*_, *INC, INF, MRCAT*) checks the sub tree *t*_*M*_ assuming each node *n* is assigned a fixed incubation time interval *inc*_*n*_ ∊ *INC* and a fixed infectious time interval *inf*_*n*_ ∊ *INF* and returns 1 if the constraints are met and 0 otherwise. The final probability calculation is the set of timing intervals that maximize the product of the individual timing probabilities over all nodes:

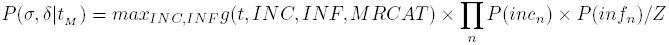

A similar procedure is applied for the linear transmission chain case. Here the algorithm starts with the observed source node and propagates candidate infection timing intervals forward to the next recipient node (rather than backwards to the parent node) until the final observable node is reached and the set of possible infection timing intervals is compared with terminal node timing to determine if a candidate timing interval supports the proposed transmission chain hypothesis.

The key drawback to the method is computational cost. There are 2^*d*+1^ timing assignments that must be considered for each combination of timing intervals. If the number of possible distinct values for *inc*_*n*_ and *inf*_*n*_ is *x* the number of timing interval combinations becomes *x*^*M*+1^. Thus, there is a practical limit to the maximal value for *M* and *x*. Measuring the potential linkages for distantly related host pairs can be determined by viewing the estimated probability values of linkage up to the maximum M and when all probability values are low, M can be inferred to be too small to allow transmission inference.

### Testing

To evaluate the potential strength of inference from combining sequence and timing data, a simulation was used to generate genetic sequences from pre-defined outbreak conditions. The simulation uses a generative stochastic process for *P* (*σ*, *δ*|*t*_*M*=1_) and takes the following parameters as input:

- A tree of infected hosts and their respective transmission links.
- Two probability density functions *P*_*inc*_ and *P*_*inf*_ determining the length of time intervals *inc*_*n*_ and *inf*_*n*_ respectively for each node *n* in a outbreak tree.
- A transmission probability density function *P*_*trans*_ determining the probability of transmitting the virus from host A to host B at time t within time interval *inf*_*A*_.
- A sample probability density function *P*_*sample*_ determining when a sequence sample is collected within time interval *inf*_*A*_ for host A.
- A per day mutation rate and an initial source index sequence.

A single run simulates an outbreak by propagating sequences across the pre-specified outbreak tree. The index case node *A*’s initial infection time is set to 0 (e.g. *E*_*A*_ = 0) and the initial infectious time *I*_*A*_ and recovery time *R*_*A*_ are determined by values drawn from the *P*_*inc*_ and *P*_*inf*_ distributions. Secondary infected host B’s initial infection time *E*_*B*_ is determined by drawing from the *P*_*trans*_ distribution to fix the time within the infectious interval from *I*_*A*_ to *R*_*A*_ that the actual transmission event takes place. The index sequence evolves daily according to the per day mutation rate with distinct sequence stored for each host *n* for each day from *E*_*n*_ to *R*_*n*_. When host *A* transmits the virus to host *B* at time *t*, the sequence in host A at time t becomes the seed sequence for host B. A single sequence is used to represent each host node, the chosen sampled sequence is determined by *P*_*sample*_, which gives a discrete time point within the infectious time interval from *I*_*n*_ to *R*_*n*_ in host node n. Sampled sequences are assumed to be taken following presentation of symptoms in the infected individual. This simulation approach is similar to a related effort to use an epidemic simulation model to construct large scale seasonal outbreaks of measles with viral genome sequence data [24]. Simulations were used to evaluate different viral sampling strategies for their proficiency in reconstructing viral population histories. A key difference with the work presented here is the focus on recovering individual host transmission chains rather than homogeneous host population infection models [25].

Using the simulated sampled sequences from host nodes, phylogenetic trees were generated with BEAST running 1,000,000 iterations with 100,000 initial runs discarded and evaluating 10,000 sampled trees. The HKY nucleotide substitution model was used with fixed variation, and a relaxed molecular clock [26] with exponential distribution and a fixed population size coalescent model. The genetic distance between two sequences was taken to be the sum of the absolute time difference between two leaves in the tree and the most recent common ancestor. The final distance value is the weighted average over all available inferred trees. The same set of most recent common ancestor times was used as input to the method for integrating time with genetic data. For the test results reported below, the probability distributions for the incubation time and infectious duration time were taken to be uniform across a pre-determined range described below. The simulator reflects reported features from the 2001 UK FMD outbreak described in [21]. Infected hosts here do not reflect individual animals but rather infected farms. An average incubation time of 5 days is assumed with a variation between 2 and 14 days determined by a gamma distribution. The infectious time interval varied from 1 to 10 days drawn from a distribution model on previously described time intervals given by [21]. The objective was to evaluate the impact of combining the genetic data with timing information and apply the evaluation methods to the original 2001 UK FMD outbreak data to assess the confidence of the associated transmission links predicted using similar evidence but using the new inference approach. A per day mutation rate of 2.226 × 10^−5^ is used following a previously published report [10].

Figure 6 shows the tree used for the simulation. The tree is based on a subset of the 2001 outbreak given in [21]. The nodes labeled with two character codes were added to the published tree to connect nodes thought to be linked indirectly such that the total time duration distribution generated from repeated simulations covered the same time frame of 110 days reported for the outbreak. Results are based on 50 independent outbreak simulations. In the genetic distance only based framework labeled training data is used to estimate the likelihood of observing a genetic distance under the different possible transmission hypotheses. In the modified framework, a prior belief of how the virus spreads replaces the evidence taken from a training set. To evaluate the two approaches, 5 fold cross-validation takes the 50 randomly generated outbreaks and partitions the outbreaks into 5 non overlapping sets of 10 distinct outbreaks to evaluate predicted transmission link accuracy using the remaining 40 outbreaks for training. Evaluations are repeated 5 times to report performance for all 50 outbreaks. When training data is not needed, statistics are simply collected on all 50 outbreaks in one step. Three basic prediction methods are compared - using genetic data only, using the epidemiology timing data only and using both lines of evidence together to inform the final prediction.

**Figure 6:**
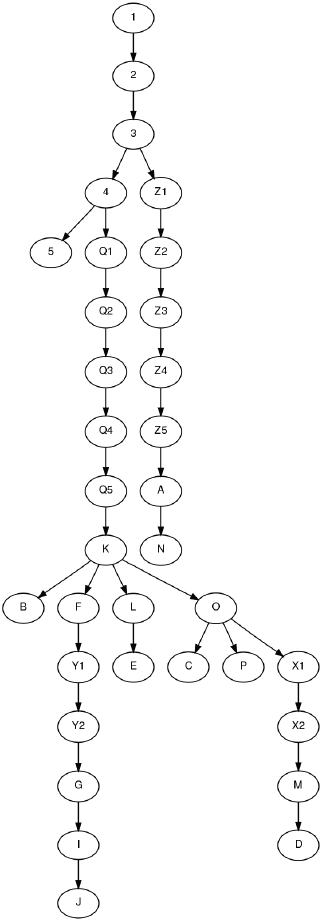
Tree used to generate simulated sequences and reflect known transmission links. Two character codes reflect hypothetical farms included in the tree and edges represent transmission relationships between infected farms.

**Figure 7:**
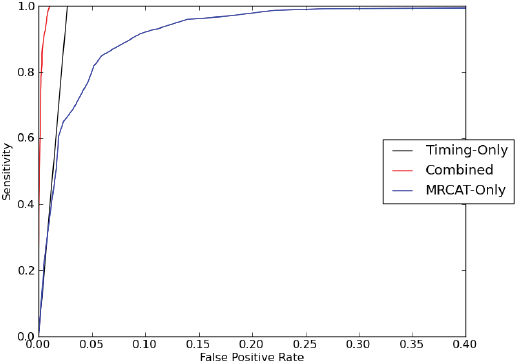
Performance with knowledge of timing. ROC curves showing the ability to correctly infer direct transmission cases (Sensitivity) versus falsely labeling indirect transmission pairs as direct transmission cases (False Positive Rate) using different confidence cut off values. Three methods are shown, using exact knowledge of timing data alone (Timing-Only), using only genetic data (MRCAT-Only) or combining the two and assuming exact knowledge of infection times.

Figure 7 shows a ROC curve for the rate of true direct transmission links identified (y-axis) versus the rate of host nodes falsely predicted to be linked (x-axis) using different prediction cutoff values. First consider prediction accuracy with near perfect knowledge of the timing events for the observed host nodes shown by the Timing-Only plot in Figure 7. The linear plot reflects the fact that the method generates a single prediction value so that above a certain cutoff either all of the true direct transmission links are predicted at the expense of falsely predicting transmission links for 3% of the other non-linked host node pairs or using a lower cutoff yields no predicted links. Recall that the calculation is meant to represent the most probable collection of incubation and infectious times consistent with the transmission hypothesis relative to the most probable collection of incubation times independent of the hypothesis. Here just a single set of incubation and infectious duration times are considered (the true time) and thus every pair of nodes with a timing pattern consistent with the predicted transmission event occurs with probability 1. Thus, the false positive rate reflects the fact that roughly 3% of node pairs that are not linked have a timing pattern consistent with the potential for direct transmission. These pairs likely refer to “siblings” or other nodes that are close by in the tree. This provides a baseline measure for quantifying the difficulty of the inference problem and shows that when exact timing knowledge is available the possible transmission inferences become highly constrained. The MRCAT-Only line shows the other end of the spectrum where there are no explicit timing constraints to prevent inference of direct transmission. The plot shows that a low confidence threshold is required to correctly identify all true direct transmission links, which shows that there is a wide overlap between the genetic distance range for both direct and indirectly linked hosts. Combining the two approaches shows that the already low false positive rate from the highly constrained timing data can be reduced still further by adding the constraints introduced by the timing data inferred from the time stamped genetic sequence data.

**Figure 8:**
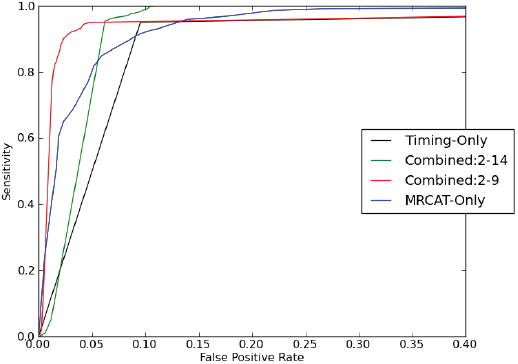
Limited timing knowledge. ROC curves showing the ability to correctly infer direct transmission cases (Sensitivity) versus falsely labeling indirect transmission pairs as direct transmission cases (False Positive Rate) using different confidence cut off values. Four methods are shown, using only timing data (Timing-Only), using only genetic data (MRCAT-Only) or combining the two assuming either a 2-14 day incubation period (Combined:2-14) or a 2-9 day incubation period (Combined:2-9).

Figure 7 illustrates how the algorithm works but assumes unrealistically precise knowledge of incubation and infectious duration times. Figure 8 shows the same comparison, but reducing the knowledge of incubation and infectious times. There are two distinct parameter settings shown in the Figure 8. One setting where every observed node is assumed to have an incubation time that could range from 2 to 9 days with equal probability and the estimated infectious duration for each node is based on the known window but assigned and uncertainty of plus or minus one day (Combined:2-9 in Figure 8). The second setting is identical except the incubation time is assumed to extend from 2 to 14 days (Combined:2-14 in Figure 8). The Timing-Only plot uses the 2 to 9 day incubation window and shows that (without using the genetic data) the range of possible timing scenarios more than triples the number of node pairs that are falsely linked by direct transmission from 3% to 10%. Combining the data sources shows improved inference with consistently higher sensitivity levels and less false positives. The results also show there are clear performance trade offs illustrating the underlying difficulty of the problem. On the one hand capping the maximal incubation time at 9 days greatly constrains the number of plausible transmission hypotheses, thus limiting the number of false positives. The trade off is that for the transmission cases where the incubation time actually exceeds 9 days the true positives can be missed, thus there is a hard upper bound on the sensitivity rate that is less than 1. Further relaxing restrictions to allow for the longer incubation times of up to 14 days allows detection of all true positives, but the number of false positives increases too. The key here seems to be the uncertainty associated with assigning a time to the most recent common ancestor. Longer incubation times open more plausible phylogenetic scenarios where two hosts are falsely linked by direct transmission but are actually linked by a shared infectious source. One potential way to strengthen inference could be to collect more sequences per farm spread over multiple time points, which in some situations may reduce the range of most likely most recent common ancestor times.

A second experiment was conducted to re-examine the transmission relationships reported by Cottom et al. on samples taken from a subset of farms infected during the 2001 United Kingdom outbreak. Although the true outbreak tree cannot be empirically determined, re-examination of this dataset proved useful in determining whether our new transmission inference method using similar information as input would lead to similar conclusions. Moreover, differences in output should highlight important differences in the two approaches and help assess the strengths and weaknesses of the different approaches.

Figure 9 shows the higher confidence inferred transmission links for the published 2001 outbreak data with whole viral genome sequence data available for each farm. An incubation window of 2 to 14 days and an estimated infectious time interval with a 1 day uncertainty was used to find the most likely transmission chain. Cottom et al. similarly apply a transmission inference algorithm to the data using a closely related approach that takes both estimated infection times and genetic sequence data as input. Cottom et al. use a statistical parsimony tree relating the collection of sequences as the basis for building putative transmission trees. The algorithm starts at the leaves of the parsimony tree and assigns the inferred common ancestor nodes to one of the farms (with a descendant leaf) as the putative source. All possible tree labellings are evaluated using an infection time likelihood score, which similar to our approach estimates likely infection times based on estimated incubation and infectious duration times. Several important differences between the two approaches emerge after examining the differences in output. One difference is that our method incorporates putative unobserved farms into transmission hypothesis tests, which help prevent over confidence in certain transmission inferences. As will be discussed in the next section, this can also form the basis for supporting a more flexible framework for evaluating more sparsely sampled outbreaks. A second difference is that multiple phylogenetic tree topologies are used in the form of multiple most recent common ancestor time estimates, which better reflect the uncertainty of phylogenetic inference. Finally, a key difference is in the information representation format. The Cottom et al. approach assigns outbreak tree nodes to phylogenetic tree nodes whereas our method assigns phylogenetic tree nodes to outbreak tree nodes, which represent variable length time. The significance of these differences is that the new framework can test many additional plausible transmission hypotheses, which would otherwise be ignored.

**Figure 9:**
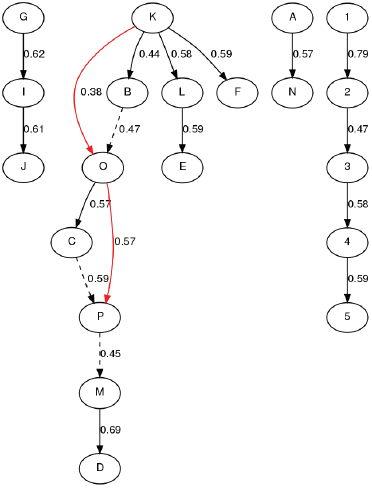
A most likely inferred transmission tree with links comparing the tree to previous reports. Red lines show novel links and dashed lines show previously reported alternative links. Each node represents an infected farm.

**Figure 10:**
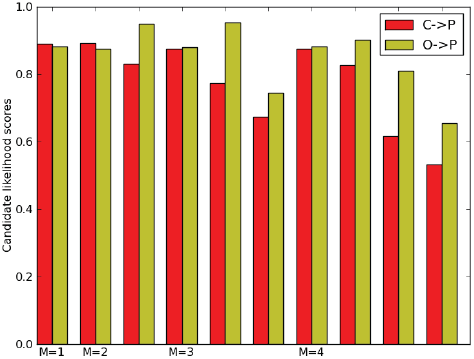
Likelihood for different transmission hypothesis for C to P and O to P, with different transmission chain lengths M.

Despite the differences, there was largely strong consensus for predictions between the two methods with all of the farms linked with high confidence in [21] predicted in the new method except the K to B link (results not shown). The contiguous edges marked in black show transmission inferences that match [21]. Linkage between K and B required increasing the infectious window beyond the initial reported estimate. To make a complete comparison the discrepancy is noted and the infectious window was increased to 4 days to accommodate the possibility of a transmission link from K to B. The two dashed edges show links made by the new inference tool not present in the most probable transmission tree reported in [21]. The two red edges show the direct transmission inferences in the most likely Cottam outbreak tree that are not present in the most probable tree reported here. Next the key differences in the two approaches are discussed with respect to differences in observed transmission inference.

- Transmission hypotheses include the potential for intermediate unobserved farm involvement to estimate confidence values.

For farm P, the new method only slightly favors an alternative source - C over O (0.57 versus 0.59) making it impossible to preclude either farm as the likely source. The favoring of C over O can be explained in part by differences in the alternative hypothesis likelihood scores. Figure 10 shows the likelihood scores for each of the individual transmission hypothesis linking C to P and O to P. The x-axis is labeled by the number of intermediate host nodes (+ 1). Adjacent unlabeled values have the same M value but a different sub tree structure. The figure shows that the direct hypothesis scores are nearly identical but the alternative hypothesis for M=2 shows that an indirect link for O to P is more plausible. Thus, the explicit scoring of alternative transmission hypotheses involving hypothetical infected farms highlight potential transmission inference ambiguity. In the case of the Cottam tree, the potential for an O to P link is not discussed, however, it appears that this possibility may be precluded due to the structure of the parsimony tree.

**Figure 11:**
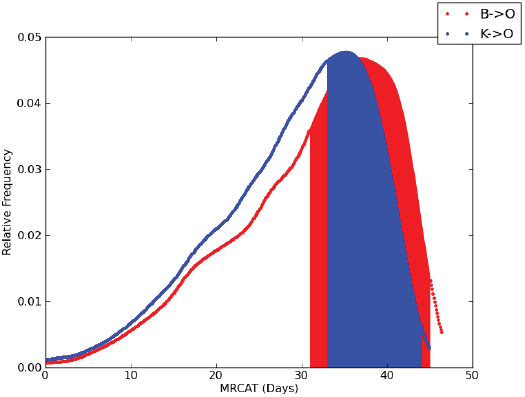
Inferred most recent common ancestor times. Range of inferred most recent common ancestor times (MRCAT) between the B,O sequence pairs (red) and the K,O sequence pairs. Shaded area under the two curves show the time intervals consistent with the respective transmission hypotheses.

- Use of multiple most recent common ancestor time estimates in addition to transmission time is included in the transmission likelihood calculation.

For farm O, the current method more clearly favors B as the source over K. Figure 11 shows weighted probability density estimates for the most recent common ancestor times for B and O (red) and K and O (blue). The red and blue shaded area under the curves show the range of predicted initial infection times consistent with the MRCAT and the direct transmission hypothesis for B to O and K to O respectively.

**Figure 12:**
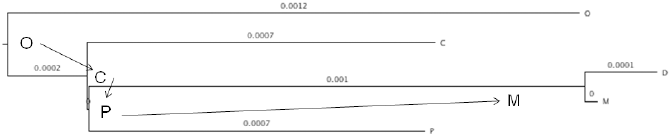
Portion of the maximum credibility tree. Phylogenetic tree leaves represent sequence taken from four farms: O, C, D, M and P. The putative transmission chain O to C to P to M is overlaid onto the phylogenetic tree, with each farm assigned to an ancestral node.

The red area exceeds the blue area indicating that the inferred MRCA times between B and O are more frequently consistent with the B to O direct transmission hypothesis than the K to O alternative. Similar to the previous case, it appears that the inferred tree used in Cottam may preclude the possibility of a K to B to O transmission path.

- Farm transmission is modeled to occur instantaneously (with respect to time) and the variable length evolution occurs within the host.

The maximum credibility tree shown in Figure 12 indicates the limitation of using the inferred ancestral nodes as the basis for reconstruction of the outbreak tree. The transmission chain O to C to P to M includes a 0 length branch from C to P that suggests a transmission from farm C to farm P with no genetic variation in the virus, which would be rejected as unlikely given minimum incubation times and the expected mutation rate. In the new method, however, since farm infections are represented by time intervals, the initial infection time of farm C can be shown to occur higher up the tree between two phylogenetic tree nodes, which would imply some mutation to occur on farm C prior to transmission to farm P. Thus, mapping the inferred phylogeny onto the outbreak tree rather than assigning outbreak nodes to phylogenetic tree nodes allows for the consideration of transmission hypotheses that would otherwise be excluded.

A strength of the Cottam et al. approach is that the transmission tree is built using all the sequences collectively by iteratively labeling the internal tree nodes rather than treating pairs of sequences independently, thus some transmission hypotheses conceivably could be more readily rejected by looking at multiple farms simultaneously. The new method shows the use of multiple tree topologies in a way that is similar to other recent applications [27] and demonstrates a natural way to incorporate multiple tree topologies in a single sequence comparison procedure. One potential improvement to the Cottam et al. approach would be to include additional candidate tree topologies for evaluation and report likelihood values that are averaged across multiple trees.

### Establishing transmission links in a sparsely sampled outbreak

While it may be possible to densely sample from an outbreak in select circumstances, in many cases only sparse sampling is possible and this is likely to be another limitation of the published Cottam et al. approach, which used dense farm sampling. Integrating phylogenetic data with infection timing data to infer more distant transmission linkages, however, is expected to be considerably more difficult. Intuitively, distinguishing between a transmission chain of length 20 from a chain of length 25 is not likely given the range of possible infection times. This is not a limitation of the method but a limitation of exploitable transmission inference knowledge. Nevertheless, a potential contribution of the method presented here is that if it is possible to relax the requirements of an exhaustive search of infection times, the framework could be used to more precisely quantify the uncertainty associated with making inherently more difficult transmission inferences and thus shed light on the conditions where more distant transmission links can be made.

To investigate this problem we introduce two random sampling steps to the algorithm, which when not implemented would otherwise preclude consideration of longer transmission chains. The first sampling step is to draw k different timing samples from the underlying incubation and infectious time duration distributions *P*_*inc*_ and *P*_*inf*_ respectively. For the results described here k was set to 50. The second random sampling step is in the tree traversal procedure illustrated in Figure 5, which is designed to determine whether there is a sequence of timing events for a particular transmission chain and a given collection of infection time durations that is consistent with the inferred most recent common ancestor time. Rather than an exhaustive search, a fixed number of random trials are taken. This approach is thus biased against inferring links requiring rarely occurring timing intervals that would be required to support the most recent common ancestor time. We note that additional sampling strategies could be implemented to make this approach more robust, however, the current modifications should be sufficient for evaluating the potential to apply this approach to larger sparsely sampled outbreaks.

For evaluation a variation on the previously described FMD outbreak simulation is used. Rather than using the smaller fixed tree topology, a tree topology was generated at random starting with an index node and randomly generating child nodes using a branching factor of 1.5, a maximal outbreak duration of 110 days and a maximal node size of 2,000. 100 trees were generated and the 40 trials that generated trees with 2,000 nodes were selected for further analysis. For each tree a pair of nodes was chosen at random to represent each path length present in the tree starting from 1 up to the maximal observed path length. To limit computational costs, only odd path lengths were evaluated.

**Figure 13:**
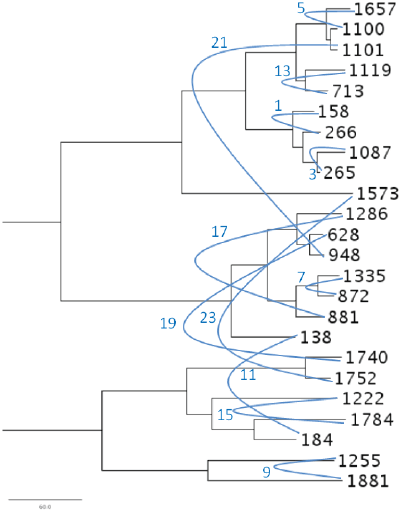
An example maximum credibility tree for a sparsely sampled simulated FMD outbreak. Tree leaves are labeled with numbers corresponding to nodes taken from a 2,000 node simulated outbreak tree. Select node pairs with known transmission relationships were evaluated. The selected node pairs are labeled by their transmission links using blue edges to link pairs and the path length marked in blue. The two sub trees are connected by a common root (not shown for image compactness).

Figure 13 shows an example consensus tree for the selected pairs taken from an outbreak chosen at random. The directed edges labeled in blue show the path lengths connecting the selected nodes with the path length labels from 1 to the maximum observed (23) for this particular outbreak tree. The tree infers evolutionary relationships that are generally consistent with what would be expected given the different transmission chain lengths observed. For example, a path length of 23 separates node 628 from 1740 and indeed the two nodes end up in two different subclades of the tree. The key question becomes can additional information be obtained by incorporating knowledge of infection times and alternative transmission hypothesis testing with the phylogenetic data to better capture the range of possible relationships among the infected farms. To answer this, we first show the likelihood evaluation for the node pairs taken from the tree in Figure 13 and then show the same likelihood evaluations averaged across all 40 randomized outbreak trials. To limit computational costs, evaluated path lengths start with 1 and 5 and continue in intervals of 5 up to 35. The likelihood value *P* (*δ|M)* is reported as the maximal likelihood value for all alternative sub trees of path length M. Figure 14 shows the value of *P* (*δ|M)* for each pair of nodes with node pairs labeled by their true transmission path length. For the sampled outbreak shown in Figure 13, with the exception of M=11, node pairs for path lengths up to 15 show *P* (*δ|M)* values that favor a specific hypothesis or a small subset of hypotheses. As the true path length increases, the average likelihood value drops and the distinction between different path lengths becomes less clear. For the node pair separated by path length of 15 for example, there are clearly higher values for inferring a path length from between 5 and 20, but there is limited distinction within this range. For the most part the framework largely reinforces what is visualized using the consensus phylogenetic tree. An interesting case, however, is the node pair (1255, 1888) linked by a transmission chain of 9, compared with node pair (713, 1119) separated by chain length 13. In the framework, both pairs of nodes have likelihood values that peak at M=5, however the alternative transmission hypotheses for linking (713, 1119) with M=10 or M=15 are much higher than they are for the (1255, 1881) case. This is an example where the relationships inferred by the tree may be misconstrued without the explicit use of a transmission inference framework. It is important to note that the transmission framework does not resolve all conflicts. For the M=11 case for node pair (138, 184), the nodes show up in two different subclades suggesting a more distant transmission relationship, than might be expected by the arrangement of other pairs in the same subclade and separated by longer transmission chains. In this case, the transmission inference framework also suggests a more distant transmission link, given the relatively uniform distribution of low *P* (*δ|M)* values.

**Figure 14:**
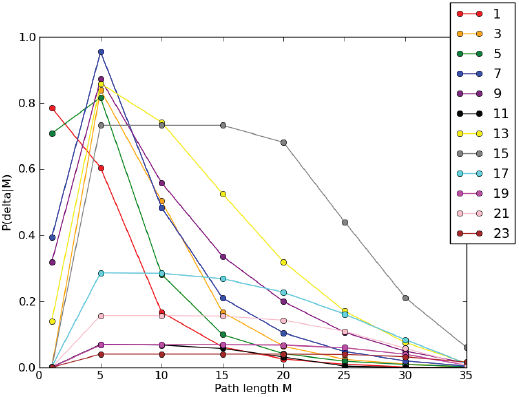
Likelihood values are given for the 12 selected transmission pairs shown in Figure 13. Each node pair is evaluated for different values of M from 1 to 35 and the true path length for each node pair is shown in the legend.

**Figure 15:**
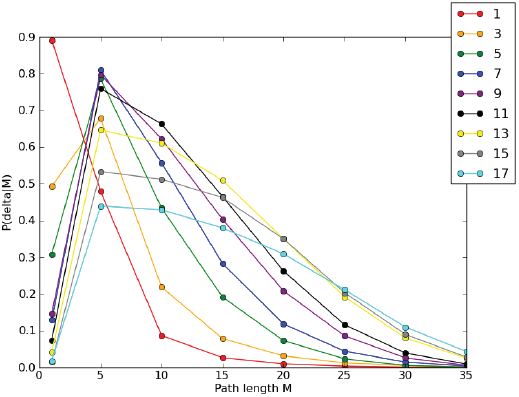
Likelihood values for all node pairs for a select path length averaged across all 40 trials. Plot shows path lengths where the average likelihood value peaks at M=5 or less.

Figure 15 and Figure 16 show *P* (*δ|M)* values averaged over the 40 random trials. For clarity the two figures are used to separate the distributions into two classes, those with peaks at *M* ≤ 5 and those that peak at *M* > 5. (Note that the maximal path length observed was 37, however, due to the 2,000 tree node limit, longer path lengths occur less frequently in the 40 trial experiment and thus path lengths with less than 40 observations are not shown.) The results show that the framework assigns a likelihood score that roughly reflects the true underlying path length. In particular, all path lengths less than 10 clearly peak at M=1 or M=5 in the figure. The results do suggest room for improvement, however, since path lengths for M=11-17 still show their maximum likelihood value at M=5. A possible reason for this is that for longer transmission chains the space of inconsistent infection times may expand at a rate that is not proportional to the increase in the number consistent infection times. Since the method uses a fixed random sample size this would favor shorter more constrained transmission hypotheses. Nevertheless, even where the likelihood value peaks at M=5, for longer transmission chains the likelihood values for *M* > 5 come increasingly closer to the peak until there is finally a peak shift at path length 19 from M=5 to M=10. It is interesting to note that as the path length increases, the peak likelihood value decreases quantifying the increasing ambiguity associated with making long transmission chain inferences. Despite this difficulty, the results indicate that in many cases it may be possible to assign a transmission relationship using a more inclusive class of transmission chain lengths. For example, nodes linked by a transmission path length less than 10 appear to be fairly distinct from those separated by path lengths greater than 20. These distinctions could prove to be of critical importance, particularly when evaluated in the context of an underlying contact network.

**Figure 16:**
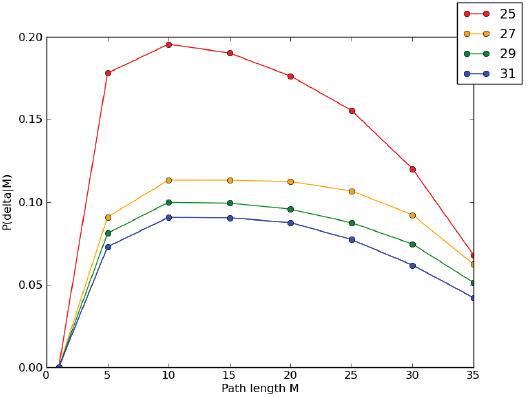
Likelihood values for node pairs connected by longer path lengths. Plot is similar to Figure 15 but shows path lengths where the average likelihood value peaks at *M >* 5.

## Discussion

It is important to note that while the presented method is designed to use prior knowledge of viral infection timing patterns, uniform probability distributions were used since an accurate estimate of the distribution is not likely in the general case. More generally since the method requires both viral evolution and epidemiology knowledge there are significant outstanding challenges not addressed in this report. The method makes the simplifying assumption of a neutral evolution model without recombination. This assumption may be appropriate for certain acute viral infection types where host immunity does not play a significant role, however, chronic infections with wider persistent circulation of multiple subtypes increase the chances to misinterpret the transmission process. Additional work is need to ensure that any violation of assumptions of evolution are detected and (ideally) corrected. The model currently lacks inter-species evolution modeling and as sequencing technology advances there will be opportunities to better measure within host viral populations [28] and the affects on the genetic features of the recipient host’s initial seeding population. A consequence of this limited knowledge is reliance on a potentially simplifying feature of viral population structure. When testing direct transmission hypotheses the most recent common ancestor between sequences sampled from the source and recipient are assumed to reside in the source host. However, with a sufficiently large and genetically diverse host population it is at least conceivable that the most recent common ancestor resides in an earlier predecessor node.

With respect to epidemiology modeling, the paper assumes there are accurate values for *P* (*M)*, which describe the prior knowledge about the relative frequency of the different types of transmission chains. There has been considerable progress in this area in recent years linking contact networks with the more traditional stochastic compartmentalized transmission models to yield estimates [11, 29] that can be inferred using knowledge gleaned from previous outbreaks. Moreover, there are other potentially useful epidemiological models that could be substituted for the SEIR approach presented here [30]. Another important area of development is the use of sequence data and a coalescent model to estimate epidemiological parameters. Recent work demonstrates the potential to estimate parameters for a SIR model from genetic sequence data [31], which could provide an important basis for specifying the epidemiological parameters for transmission inference, when limited non-genome sequence derived observation is available. A strength of the approach is to provide a framework that can readily make use of increasing amounts of knowledge for a particular outbreak. For example, awareness of infections may increase during the course of an outbreak and thus favor using two different priors for the infectious duration times, one at an early outbreak stage and another at a later stage of the outbreak. Other clinical features of infection can be taken into account either by changing prior likelihoods for alternatives in the underlying outbreak tree structure or by making adjustment to priors in the timing distributions. In some cases prior knowledge could be used to narrow the window of possible infection times and greatly strengthen the confidence of transmission inference. In other cases prior knowledge may end up increasing the window of possible infection times and thus highlight the inherent complexity of reconstructing transmission events.

## Conclusion

A new method is presented that takes as input a collection of time-stamped viral genomes and basic knowledge about the timing features of the infection, the underlying transmission network and returns confidence values that assess the probability of correctly inferring host transmission links. Simulations of an FMD outbreak show how the combined use of the epidemiology based timing constraints with the phylogenetic inferred most recent common ancestor times improve confidence in transmission inference over methods that rely on just one individual feature. The simulations show that labeled training data measuring genetic distance between direct and indirect transmission cases can be used without the additional knowledge of the features of the viral transmission, however, appropriate training data in some cases may not be available. The approach presented is expected to provide a basis for building flexible tools to more readily map inferred viral evolutionary relationships onto epidemiological models to more accurately quantify host transmission relationships.

## Acknowledgements

### Authors’ contributions

JEA and SPV conceived and designed methods. JEA implemented experiments and drafted the manuscript. JEA and SPV analyzed results.

### Acknowledgments

The authors acknowledge funding for this work from the U.S. Department of Homeland Security. This work was performed under the auspices of the U.S. Department of Energy by Lawrence Livermore National Laboratory under Contract DE-AC52-07NA27344.

